# Shared genetic contribution to type 1 and type 2 diabetes risk

**DOI:** 10.1101/285304

**Authors:** Anthony Aylward, Joshua Chiou, Mei-Lin Okino, Nikita Kadakia, Kyle J Gaulton

## Abstract

The role of shared genetic risk in the etiology of type 1 diabetes (T1D) and type 2 diabetes (T2D) and the mechanisms of these effects is unknown. In this study, we generated T1D association data of 15k samples imputed into the HRC reference panel which we compared to T2D association data of 159k samples imputed into 1000 Genomes. The effects of genetic variants on T1D and T2D risk at known loci and genome-wide were positively correlated, which we replicated using data from the UK Biobank and clinically-defined diabetes in the WTCCC. Increased risk of T1D and T2D was correlated with higher fasting insulin and fasting glucose level and decreased birth weight, among T1D- and T2D-specifc correlations, and T1D and T2D associated variants were enriched in regulatory elements for pancreatic, insulin resistance (adipose, CD19+ B cell), and developmental (CD184+ endoderm) cell types. We fine-mapped causal variants at known T1D and T2D loci and found evidence for co-localization at five signals, four of which had same direction of effect, including *CENPW* and *GLIS3.* Shared risk variants at *GLIS3* and other signals were associated with measures of islet function, while *CENPW* was associated with early growth, and we identified shared risk variants at *GLIS3* in islet accessible chromatin with allelic effects on islet regulatory activity. Our findings support shared genetic risk involving variants affecting islet function as well as insulin resistance, growth and development in the etiology of T1D and T2D.

## Introduction

Diabetes affects over 400 million individuals worldwide and contributes to substantial morbidity and mortality^1^. Type 1 diabetes (T1D) is an autoimmune disease resulting in destruction of pancreatic beta cells, whereas type 2 diabetes (T2D) is a metabolic disease of insulin resistance and beta cell dysfunction^2^. Genetics plays a major role in both forms of diabetes, where 58 risk signals have been identified for T1D^3^ and over 100 for T2D^4,5^. Roughly half of the genetic risk for T1D can be attributed to the HLA locus, and many known T1D risk loci affect immune function^2^. Conversely, the majority of known T2D risk loci appear to affect pancreatic islet and insulin resistance tissues such as adipocytes and skeletal muscle^6–10^. Outside of known loci there are many additional genetic factors influencing diabetes risk^8^. Pathophysiological links have been reported between T1D and T2D suggesting an underlying shared etiology^11,12^, but the contribution of genetic variants to this shared etiology and the underlying molecular and physiological mechanisms are unknown.

Multiple genomic loci that affect risk of both T1D and T2D have been identified. One example is the *CTRB1* locus, where risk variants are correlated with chymotrypsin expression in the pancreas and pancreatic islets and GLP-1 mediated insulin secretion^13^. Another example is *GLIS3*, a gene that causes monogenic neonatal diabetes^14^. A linkage study in non-obese diabetic (NOD) mice identified an effect of the *GLIS3* locus on T1D progression, suggesting an underlying pancreatic beta cell phenotype^12^. This study further argued that beta cell ‘fragility’ involving the unfolded protein response leading to pronounced cell death underlies shared T1D and T2D risk^15^. However, the specific causal variants at shared risk loci, including whether the signals are the same or distinct, and the mechanisms of how they alter genomic and cellular functions to influence disease risk are unknown. Furthermore, shared loci appear to have both opposite (*CTRB1*) and same (*GLIS3*) direction of effect on T1D and T2D risk, and thus the broader relationship between genetic effects on T1D and T2D is unclear.

Genome-wide association data of variant genotypes imputed into comprehensive reference panels enables understanding broad relationships to other traits and functional annotations^16–18^. In addition, these data enable fine-mapping of causal variants and mechanisms underlying diabetes risk at specific loci^8^. Previous fine-mapping studies of T1D and T2D loci resolved sets of causal variants at many risk signals and annotations enriched in these causal variant sets^3,19^. These studies revealed that the majority of risk signals for diabetes map in regulatory elements active in specific cell-types and thus likely affect gene regulation in these cells^3–8,19^. Projects such as ENCODE and the NIH Epigenome Roadmap have annotated regulatory elements in hundreds of human cells and tissues^20,21^, while other studies have provided detailed regulatory maps of specific tissues such as islets and adipocytes^6,22^. Epigenomic annotations broadly enriched for disease signals can further be used to prioritize potential functions of causal variants overlapping these annotations for experimental validation^19^.

Here, we studied genetic risk of T1D and T2D using comprehensive genome-wide association data for both traits. We identified positive correlations both genome-wide and at known loci between variant effects on T1D and T2D risk. Increased risk of T1D and T2D was correlated with higher fasting insulin and glucose level and decreased birth weight, among other traits, and variants with T1D and T2D association were enriched in pancreatic islet, adipocyte, CD19+ B cell, and CD184+ endoderm regulatory elements. We identified evidence of co-localized signals for T1D and T2D at five loci, four of which had the same direction of effect. Shared signals at *GLIS3* and other loci were associated with quantitative measures of beta cell function, while *CENPW* was associated with early growth phenotypes. We fine-mapped casual variants at shared signals and identified variants at *GLIS3* in islet accessible chromatin with allelic effects on enhancer activity. Together our results provide evidence for shared risk underlying T1D and T2D involving variants with effects on pancreatic islets and well as insulin resistance, growth, and development.

## Results

We generated genome-wide association data for T1D using publicly-available genotype data of T1D case and control samples of European ancestry (**see Methods, Figure S1**). We imputed genotypes from each study into 39M variants in the Haplotype Reference Consortium (HRC) panel^23^. Imputed genotypes passing quality filters (r^2^>.3) were tested for T1D association separately for different genotyping platforms using firth-biased regression including sex and the top 3 principal components as covariates. We then performed inverse variance weighted metaanalysis to combine results. We retained imputed variants tested in all samples with minor allele frequency (MAF) > .005, resulting in 8.5M variants. As expected, given comparable sample size to previous studies, variants with genome-wide significant association mapped to known loci (**Figure S1**).

We then determined the relationship between variant effects on T1D and T2D risk by comparing T1D association statistics with T2D association from the DIAGRAM consortium^4^. We first determined shared effects among variants at known risk loci for both traits excluding the MHC locus. There was an enrichment of nominal T1D association (P<.05) among 93 known T2D index variants relative to matched background variants (obs=19.1%, exp=7.8%, binomial P=3.2×10^-4^) (**Figure 1A, Table S1**). T2D index variants were also enriched for concordant direction of effect on T1D (57/94, binomial P=.037), including among those with nominal T1D association (T1D P<.05) (14/18, binomial P=.031) (**Figure 1B, Table S1**). We found significant directional concordance among the 14 variants with both nominal T1D association and same direction of effect on T2D using summary data from UK Biobank (UKBB) (12/14, binomial P=.013). Despite a net sharing in effects, several T2D loci had opposite effects on T1D risk including *CTRB1* and *TCF7L2* (**Figure 1B**). Conversely, there was less evidence for enrichment of nominal T2D association (obs=12.2%, exp=7.3%, binomial P=.19) or concordant direction of effect (28/57, binomial P=1) among 57 known T1D variants (**Figure 1A, Table S2**).

**Figure 1.**
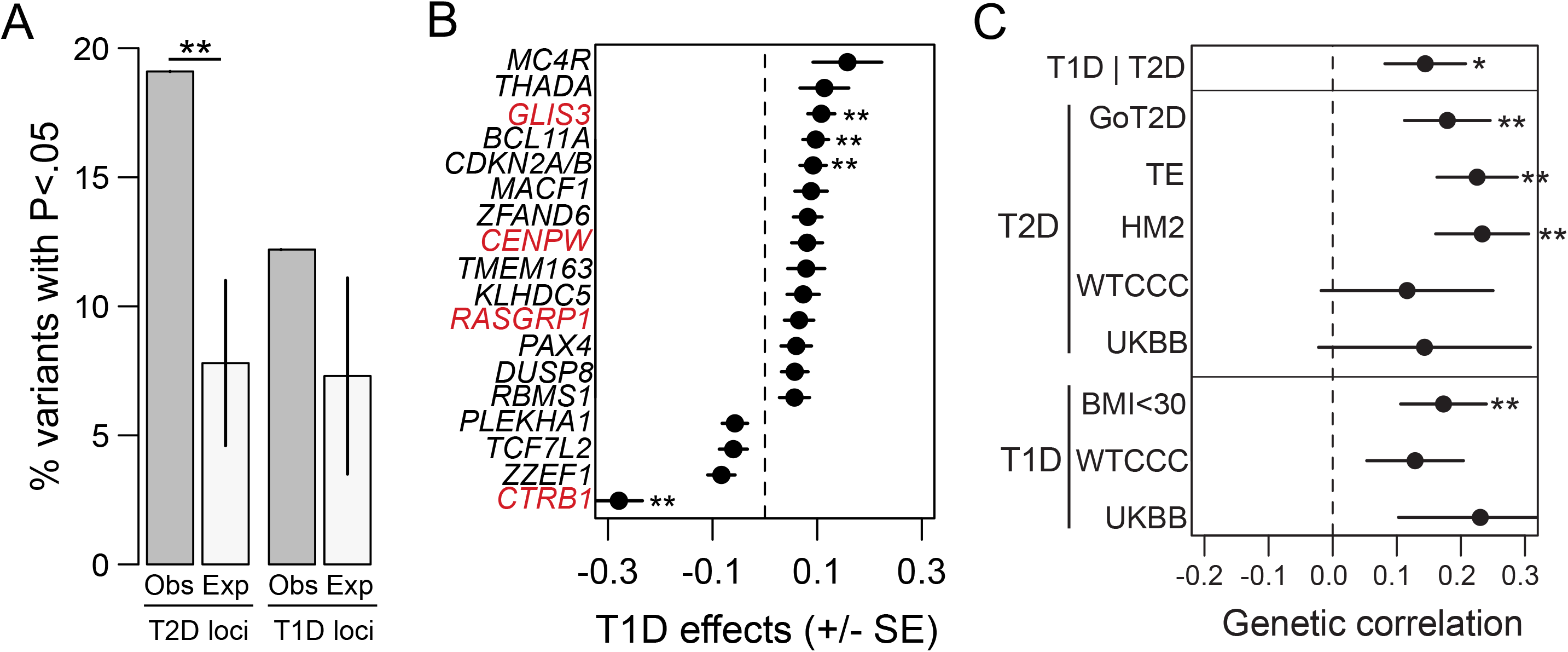
Shared effects of genetic variants on T1D and T2D risk. (A) Known T2D risk variants are significantly enriched for nominal T1D association (P<.05), whereas known T1D risk variants do not show evidence for enrichment of nominal T2D association. **P<.001 (B) Known T2D risk variants with nominal T1D association have concordant direction of effect on T1D risk (14/18, red=known T1D locus; **index variant T1D P<5×10^-4^). Values are T1D effect size and SE. (C) Variants genome-wide have correlated effects on T1D and T2D risk using multiple datasets for each disease (UKBB – UK Biobank, WTCCC – Wellcome Trust Case Control Consortium, T2D TE – Mahajan et al, T2D HM2 – Morris et al 2012, T2D GoT2D – Fuchsberger et al 2016, T1D BMI<30 – T1D association data removing obese case samples). Values are heritability estimates and SE.

We then determined the correlation between variant effects genome-wide on T1D and T2D risk. In these analyses, we used LD-score regression on the set of HapMap3 variants common to T1D and T2D association datasets (**see Methods**). We observed evidence for a positive correlation in the effects of variants genome-wide on T1D and T2D risk (R_g_=.14) (**Figure 1C**). A positive correlation remained when performing these analyses using summary data of T1D and T2D from the UK Biobank (T1D/T2D-UKBB R_g_=.12, T1D-UKBB/T2D R_g_=.23) (**Figure 1C**). We also identified positive correlation with T1D risk when using T2D association data imputed from different reference panels (GoT2D, HM2) (R_g_=.18, R_g_=.23) and from trans-ethnic cohorts (R_g_=.22) (**Figure 1C**). To limit the potential effects of misdiagnosed diabetes on these results, we first generated association data using clinical definitions of T1D and T2D in the WTCCC and observed a positive correlation when using either T1D or T2D WTCCC dataset (T2D-WTCCC R_g_=.14; T1D-WTCCC R_g_=.13) (**see Methods**). Second, we removed obese (BMI>30) samples from T1D cohorts and the positive correlation with T2D remained (R_g_=.14) (**Figure 1C**). These results demonstrate evidence for correlated effects of variants genome-wide on risk of T1D and T2D.

Given evidence for a positive correlation in variant effects on T1D and T2D, we sought to understand potential mechanisms underlying the shared effects. We first determined the correlation between T1D and T2D risk and relevant traits using LD score regression^24–27^. For T2D, there was a significant correlation between T2D risk and increased HbA1C level (R_g_=.64, P=3.1×10^-15^), fasting glucose level (R_g_=.57, P=4.2×10^-11^), fasting insulin level (R_g_=.48, P=2.9×10^-9^), HOMA-IR (R_g_=.55, P=1.9×10^-7^), and body-mass index (BMI) (R_g_=.48, P=3.9×10^-36^), and decreased birth weight (R_g_=-.28, P=1.2×10^-8^) (**Figure 2A**). There was also evidence for a correlation between T2D risk and increased proinsulin level (R_g_=.22, P=.057) and male pubertal size (R_g_=. 12, P=.14) although these estimates were not significant. For T1D, we observed a correlation between T1D risk and increased fasting proinsulin (R_g_=.23, P=.034) and fasting insulin level (R_g_=.17, P=.047) (**Figure 2A**). We also observed evidence for a correlation between T1D risk and decreased birth weight (R_g_=-.09, P=.10), increased male pubertal size (R_g_=.18, P=.11), and increased fasting glucose level (R_g_=.07, P=.32) although these estimates were not significant. We did not observe correlation between T1D and BMI (R_g_=-0.02, P=.52) or childhood obesity (R_g_=-0.02, P=.75), the latter previously identified as an instrumental variable for T1D risk^28^.

**Figure 2.**
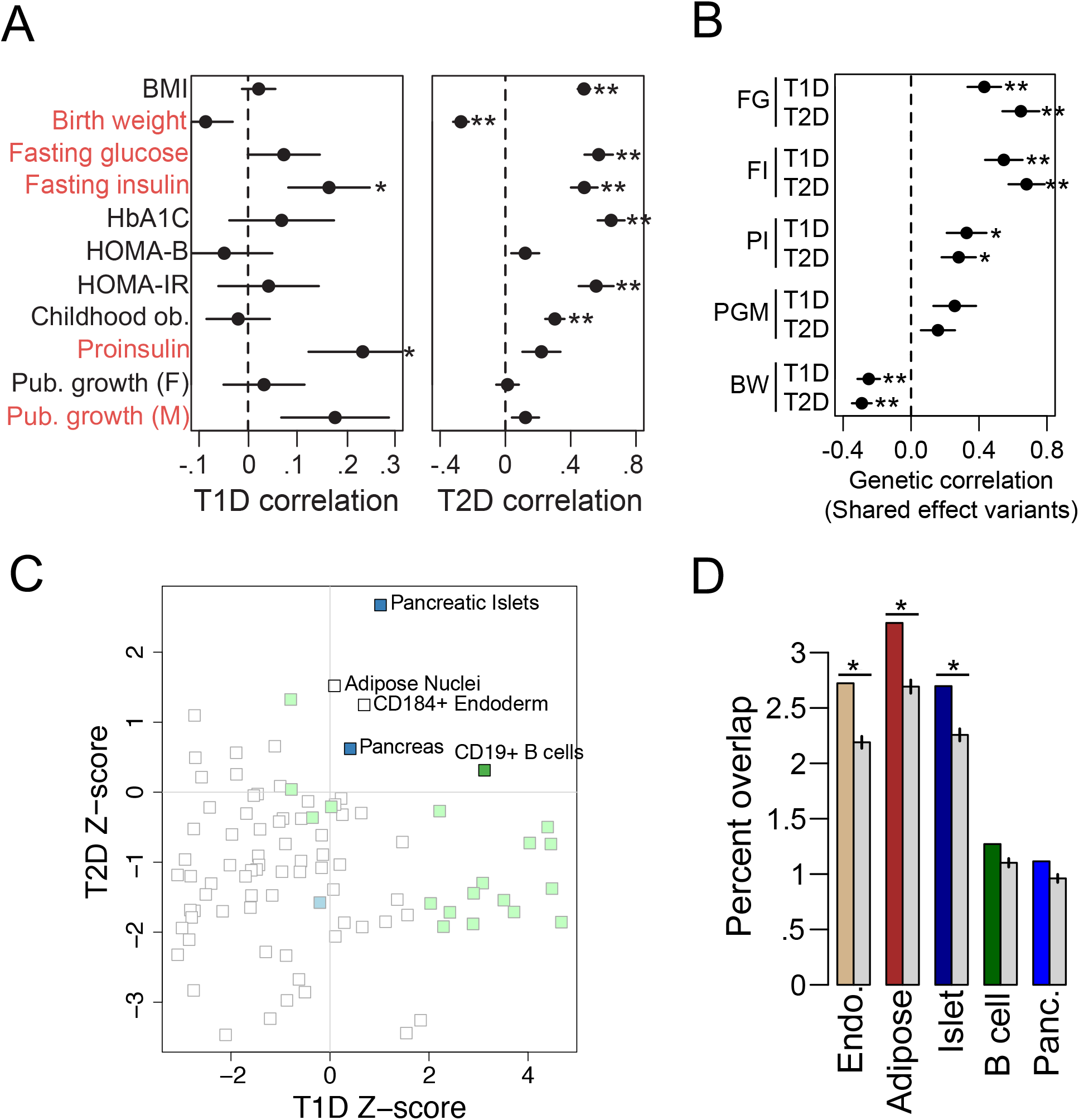
Mechanisms of variant effects on T1D and T2D risk. (A) Increased T1D risk (left) is correlated with increased fasting insulin level and proinsulin level (*P<.05), in addition to increased male pubertal growth and fasting glucose level, and decreased birth weight; Increased T2D risk (right) is correlated with increased HbA1C, fasting glucose, fasting insulin, HOMA-IR, BMI and childhood obesity, and decreased birth weight (**P<1×10^-4^). Values are heritability estimates and SE. (C) Variants with same direction of effect on T1D and T2D risk have stronger correlation with increased fasting insulin, glucose and proinsulin level, and decreased birth weight. (**P<1×10^-4^, *P<.05). Values are heritability estimates and SE. (D) Variants with T1D and T2D association are enriched for pancreatic islet, adipose, CD19+ B cell, and CD184+ endoderm regulatory sites. (blue = pancreatic, green = immune). (E) Variants with both nominal association (P<.05) and shared direction of effect on T1D and T2D risk are significantly enriched in endoderm, islet and adipose regulatory sites. (*P<.05). Values are percent overlap and CI.

We determined the extent to which traits correlated with both T1D and T2D risk might be driven through variants with shared effects on T1D and T2D. From genome-wide association data for T1D and T2D, we extracted variants with the same direction of effect and tested these variants for correlation to each trait using LD score regression. For both T1D and T2D, we observed stronger correlations with increased fasting glucose level (T1D shared R_g_=.43, T2D shared R_g_=.65), increased fasting insulin level (T1D shared R_g_=.55, T2D shared R_g_=.68), and decreased birth weight (T1D shared R_g_=.25, T2D shared R_g_=.29) among variants with same direction of effect (**Figure 2B**). We observed less evidence for pronounced correlation between shared effect T1D and T2D variants and fasting proinsulin level (T1D shared R_g_=.33, T2D shared R_g_=.28), and male pubertal growth (T1D shared R_g_=.26, T2D shared R_g_=.16) (**Figure 2B**).

We next determined functional annotations enriched for T1D and T2D associated variants. We used annotations of active enhancer and promoter elements in 98 cell types from the Epigenome roadmap project^21^ and annotations of protein-coding gene exons and UTRs from GENCODE^29^. We tested for enrichment of each annotation for T1D and T2D risk using stratified LD score regression^17^. There was evidence for positive enrichment genome-wide of both T1D and T2D association for variants in pancreatic islet (T1D Z=1.02, T2D Z=2.67), adipose nuclei (T1D Z=.09, T2D Z=1.52), CD19+ B cell (T1D Z=3.12, T2D Z=.31), CD184+ endoderm (T1D Z=.62, T2D Z=1.25), and pancreas (T1D Z=.41, T2D Z=.62) regulatory elements (**Figure 2C**). We also observed enrichments specific to each trait, most notably T1D association for immune regulatory elements such as T cell (Z=4.67) and fetal thymus (Z=1.83) (**Table S4**).

Given enrichment of multiple cell-types for both T1D and T2D association, we next tested to what extent these effects were driven through variants with same direction of effect on T1D and T2D. We obtained LD-pruned variants nominally associated (P<.05) with both T1D and T2D and with same direction of effect and tested for enrichment of overlap with each annotation compared to random sets of matched variants (**see Methods**). We observed significant enrichment of overlap with CD184+ endoderm (Fisher’s P=.017), adipose nuclei (P=.018) and pancreatic islet (P=.040) regulatory sites (**Figure 2D**). We next repeated these analyses instead using variants with opposite direction of effect on T1D and T2D. We observed significant overlap of opposite effect variants with CD184+ endoderm (P=.031) and pancreatic islet regulatory elements (P=.020), suggesting that these cell-types are enriched in variants with both shared and opposite effects on T1D and T2D.

We next used association data to fine-map specific causal variants influencing T1D and T2D. For T2D we compiled fine-mapping data of 93 signals from previous studies (**see Methods**). As fine-mapping data for all known T1D loci have not been previously reported, we used T1D association statistics to fine-map 57 T1D risk signals excluding the MHC region. At each locus, we considered the index variant for the locus and all variants in at least low LD (r^2^>.1). We then used a Bayesian approach to calculate the posterior causal probability (PPA) for each variant, and ‘credible sets’ of variants explaining 99% of the total PPA (**see Methods, Figure 3A, Table S5**). T1D credible sets contained a median of 66 variants, and 15 loci had 25 or fewer credible set variants. We compared fine-mapping for 34 loci common to our data and Immunochip fine-mapping^3^, and found a strong correlation between T1D association for credible set variants (Pearson r=.93). Credible set sizes at these 34 loci were larger in our data than for Immunochip (median=37, Immunochip median=31), likely reflecting increased variant density. We also identified high probability variants not covered in Immunochip credible sets for example at *CTSH* (rs12592898, PPA=.19).

**Figure 3.**
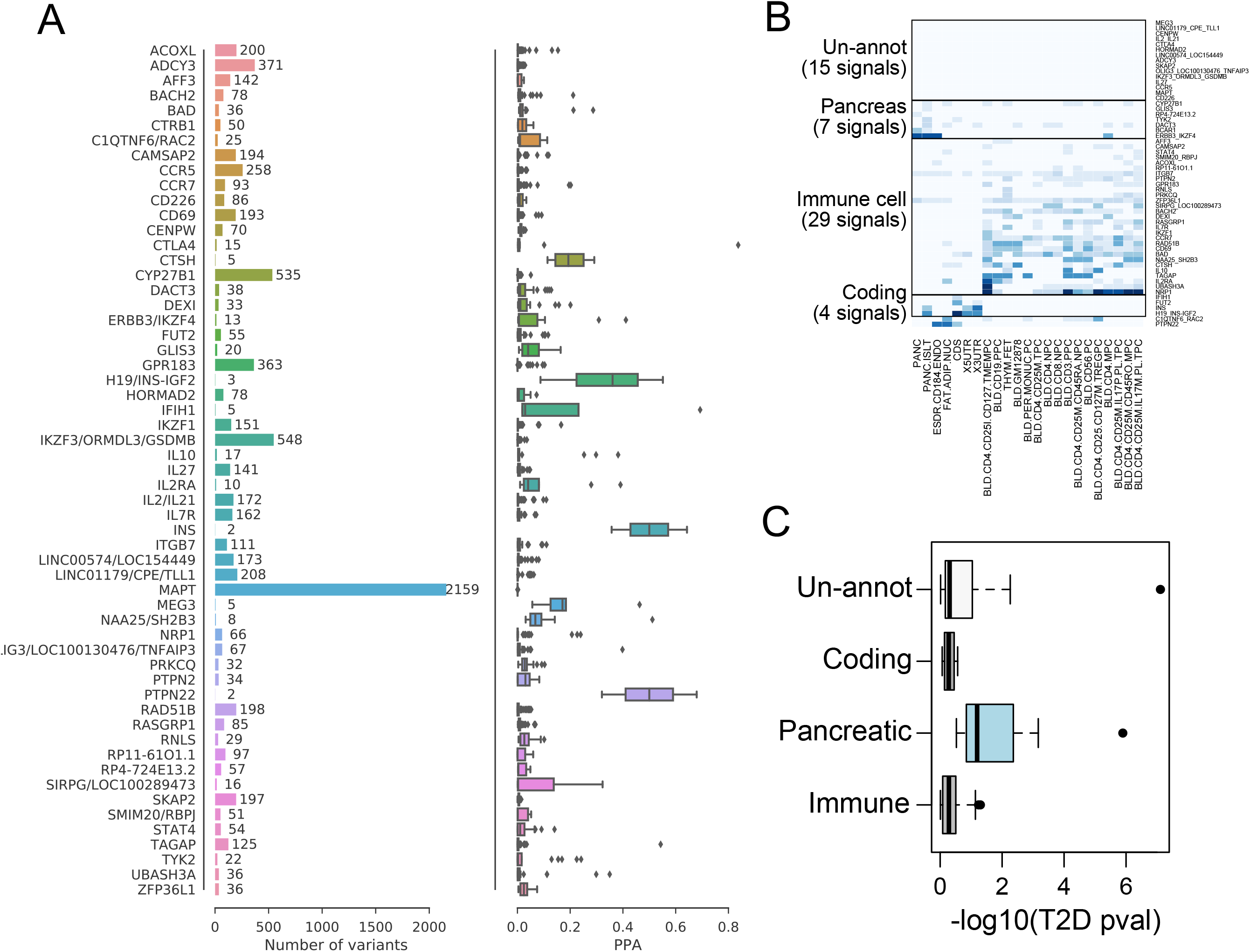
Fine-mapping and functional annotation of known T1D loci. (A) Fine-mapping of causal variant sets at 57 known T1D risk signals. (left) number of 99% credible set variants at each locus and (right) causal probabilities (PPA) of credible set variants at each locus. (B) Cumulative PPA values of 57 T1D signals in cell-type regulatory site and coding annotations. T1D signals mapped into four primary groups including immune cell regulatory sites (31 signals), pancreas regulatory sites (6 signals), and coding exons (4 signals). (C) T1D signals within different groups had distinct patterns of association with T2D, where T1D pancreas signals had the strongest evidence for T2D association.

Given fine-mapping of known T1D and T2D signals, we next determined genomic annotations of candidate causal variants at these signals. For each signal, we calculated the cumulative PPA of variants overlapping T1D/T2D enriched annotations including pancreas, adipose, endoderm and immune cell regulatory elements as well as protein-coding exons. We then grouped signals based on the resulting cumulative PPA values for each annotation (**see Methods**). For T1D, signals mapped into distinct groups of immune cell regulatory elements (31 signals), pancreas regulatory elements (6 signals), and coding exons (4 signals) as well as 15 un-annotated signals (**Figure 3B**). For T2D, signals also mapped into distinct groups including pancreas regulatory elements (21 signals), adipose regulatory elements (15 signals), and coding exons (4 signals) (**Figure S2**). T1D pancreas signals were associated with T2D risk (median -log10(P)=1.37), whereas other T1D groups did not show evidence for T2D association (**Figure 3C**). Among T1D signals in the pancreas group were those with known T2D association such as *GLIS3* and *CTRB1*, as well as others with nominal T2D association such as *ERBB3.*

Several loci have been reported to influence risk of both T1D and T2D, but whether risk signals have shared or distinct causal variants is unknown. We cataloged 144 loci with known association to either form of diabetes and tested for shared causal variants using Bayesian co-localization (see **Methods, Table S6**). There was co-localization of risk signals (P_shared_>.50) at three known T1D and T2D loci *CENPW* (P_shared_=.88), *CTRB1* (P_shared_=.88), and *GLIS3* (P_shared_=.62) as well as evidence for putative co-localization of signals at known T2D loci *BCL11A* (P_shared_=.73) and *THADA* (P_shared_=.68) (**Figure 4A**). All shared risk signals except for *CTRB1* had the same direction of effect on T1D and T2D risk. At *RASGRP1*, which has reported association to both T1D and T2D, we found no evidence for either state (P_distinct_=.03, P_shared_=.02) (**Table S5**). At several loci including *MTMR3* and *ZMIZ1*, there was evidence for two distinct T1D and T2D signals (P_distinct_>.5) (**Figure 4A**). We fine-mapped causal variants at co-localized signals by combining T1D and T2D evidence (**see Methods**). There was a reduction in credible set size at shared loci, including fewer than 10 variants at *GLIS3* (9 vars) and *CTRB1* (8 vars) (**Figure 4B, Figure S3, Table S7**). We further confirmed evidence (CLPP>.01) for shared causal variants at the *GLIS3* and *CTRB1* signals using eCAVIAR (**see Methods, Figure S3, Table S7**).

**Figure 4.**
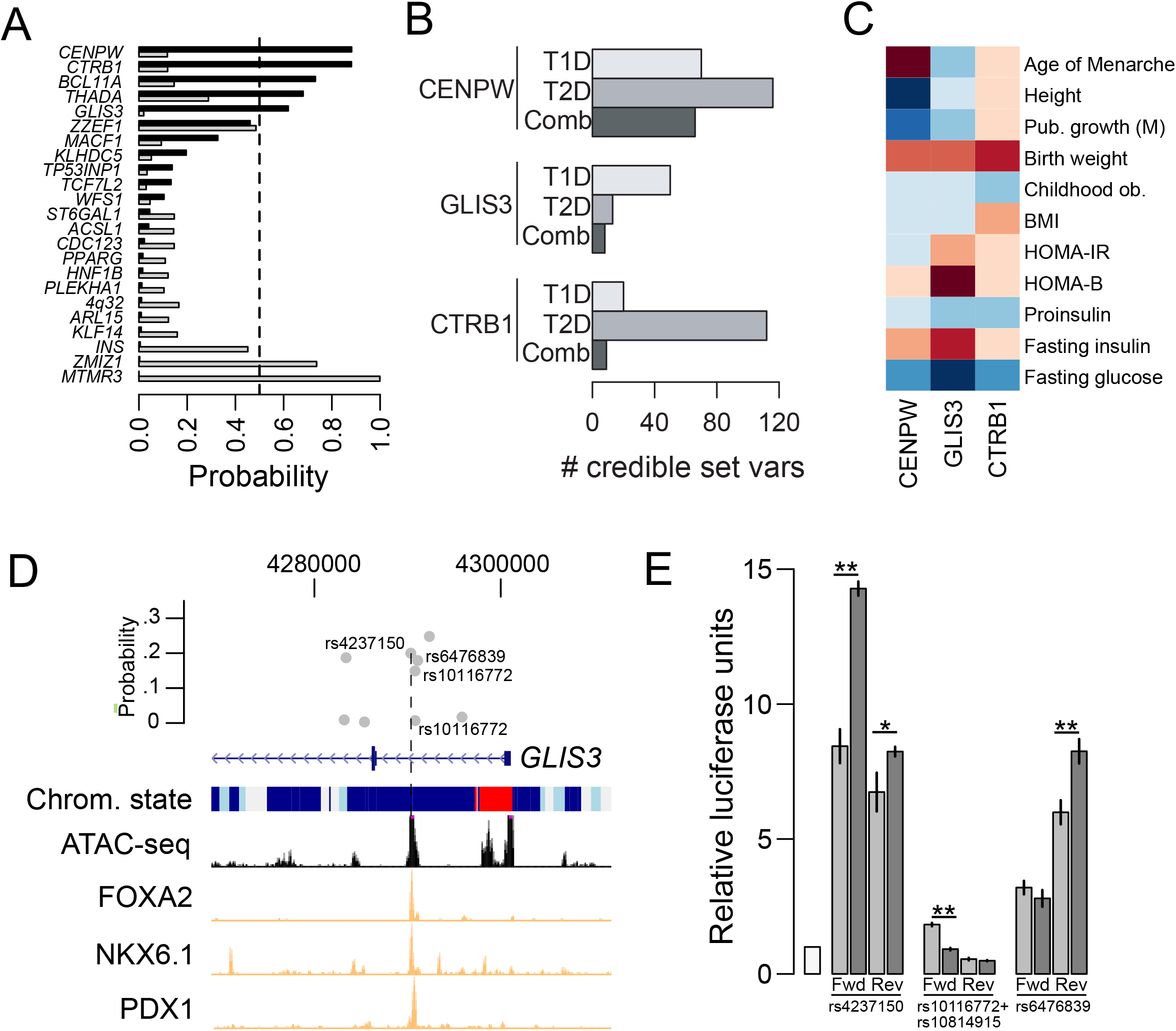
Shared T1D and T2D risk variants affect islet regulatory activity. (A) Five loci have evidence for a shared signal (P_shared_>.50) influencing both T1D and T2D risk, and two have evidence for distinct signals (P_distinct_>.50) (dark grey = P_shared_, grey = P_distinct_) (C) Number of 99% credible set variants at shared T1D and T2D risk loci. After combining T1D and T2D evidence the *GLIS3* and *CTRB1* signals have <10 variants. (C) Quantitative trait association at shared T1D and T2D signals. Values represent signed z-scores for the risk allele of the most likely causal variant (blue = positive, red = negative). For *CTRB1* z-scores are signed to the T2D risk allele. (D) Shared risk variants rs4237150, rs10116772, and rs10814915 at *GLIS3* are in islet active enhancer and accessible chromatin, and rs4237150 is also in islet TF ChIP-seq (states: dark blue = active enhancer, light blue = weak enhancer, red = active promoter) (E) Variants at *GLIS3* have allelic effects on enhancer activity in islet cells. Values are mean and SD. (N=3; *P<.05, ** P<.01).

To understand mechanisms of how the shared T1D and T2D signals influence diabetes risk, we examined quantitative trait associations at shared signals^24,30–32^. At *GLIS3*, risk alleles were associated with increased fasting glucose level (rs10758593 Z=4.51) and decreased HOMA-B (Z=-4.54) as well as decreased birth weight (Z=-2.27) (**Figure 4C**). At *CTRB1*, risk alleles for T2D were nominally associated with higher fasting glucose (rs8056814 Z=2.27) and decreased birth weight (Z=-3.78). At *CENPW*, risk alleles were also nominally associated with higher fasting glucose (rs4565329 Z=2.32) and decreased birth weight (Z=2.97), as well as increased male pubertal size (Z=3.14), height (Z=13), and earlier age of menarche (Z=-8.9). Among putative shared signals, variants at *THADA* were associated with increased fasting glucose level (Z=3.65) and decreased HOMA-B (Z=-4.23).

Multiple shared T1D and T2D signals likely affect beta cell function, and thus we annotated variants in islet regulatory sites at these signals. We used accessible chromatin sites merged from ATAC-seq in six islet samples^33,34^ (**Table S8**), chromatin states created from islet histone modification ChIP-seq data^6,35^, islet transcription factor (TF) ChIP-seq sites^6^, and TF footprints generated in islet ATAC-seq using CENTIPEDE^33^ (**see Methods**). At *GLIS3*, rs4237150 (PPA=.20), rs10116772 (PPA=.15) and rs10814915 (PPA=.007) mapped in islet accessible chromatin, active enhancer, and disrupted TF footprints, as well as islet TF ChIP-seq for rs4237150 (**Figure 4D, Table S7**). At *CTRB1*, rs8056814 (PPA=.91) also mapped in islet accessible chromatin, active enhancer and disrupted TF footprints (**Figure S4, Table S7**).

We tested these shared variants at *GLIS3* and *CTRB1*, and another nearby *GLIS3* candidate variant rs6476839, for effects on islet regulatory activity. We cloned sequence surrounding variant alleles into reporter vectors in both forward and reverse orientations, and transfected constructs into the islet cell line MIN6. As rs10116772 and rs10814915 were within 3bp, we cloned these variants in the same construct. At *GLIS3*, there was a significant allelic difference in enhancer activity in both orientations for rs4237150 (Two-sided t-test Fwd P=1.2×10^-4;^ Rev P=.024), as well as evidence in one orientation only for the rs10116772+rs10814915 and rs6476839 constructs (**Figure 4E**). We further identified evidence for allelic imbalance in islet ChIP-seq reads from samples estimated to be heterozygous for these *GLIS3* variants (see Methods; **Figure S5**). At *CTRB1*, we observed significant allelic difference in repressor activity for rs8056814 (Fwd P=.017^;^ Rev P=6.7×10^-4^; **Figure S4**).

## Discussion

Comparison of variant effects on T1D and T2D genome-wide, across known loci, and at individual loci provide evidence for shared genetic risk underlying the two major forms of diabetes. A recent study determined that a subset of patients with later-onset T1D are misdiagnosed with T2D^36^. This is unlikely to explain a positive correlation between T1D and T2D given that we observed no enrichment of T2D association or concordance in effect direction among known T1D variants, even among large effect T1D variants, and the correlation remained when using clinically defined T2D in the WTCCC with no T1D relatives, negative anti-GAD, and >1 year from diagnosis to insulin treatment. Misdiagnosis of T2D as T1D is also an unlikely explanation of the positive correlation as it remains when using clinically defined T1D in the WTCCC with onset <17, insulin treatment from diagnosis for >6 months, and no monogenic diabetes, or when removing obese individuals from T1D cohorts. Furthermore, we found little evidence for directional consistency among largest effect T2D variants.

Reports have argued that islet dysfunction underlies shared etiology of T1D and T2D^12^. Our findings support a role for shared variants at *GLIS3* in islet function, where risk alleles were associated with increased fasting glucose level and decreased beta cell function. In addition, multiple shared risk variants at *GLIS3* had allelic effects on islet enhancer activity and one was predicted to bind the glucocorticoid receptor, which is involved in diabetes-relevant inflammatory response^37^. The mechanism of how these variants influence diabetes risk through regulation of *GLIS3* and/or other genes in islets remains to be uncovered. Putative shared risk signals at *THADA* were associated with increased glucose level and decreased beta cell function, in line with a previous report^38^, and variants at *BCL11A* have been reported to affect beta cell function^38^. Candidate genes at these loci are involved in apoptotic and stress-related processes^39,40^ and therefore altered activity could contribute to a fragile beta cell phenotype. Genome-wide, T1D and T2D associated variants were enriched in islet regulatory elements and correlated with increased fasting glucose level. Given the role of islet stress response in shared risk, studies mapping the islet epigenome and gene expression in diabetogenic stress conditions will help uncover additional relevant islet regulatory programs.

Shared variants at the *CTRB1* locus have opposite effects on T2D and T1D risk and have allelic effects on islet regulatory activity, in line with a previous report correlating risk variants with *CTRB1/2* expression in pancreas and pancreatic islets^13^. The variant affects a site with apparent repressive activity in islets. Other loci have evidence for opposite effects on T1D and T2D such as *TCF7L2*, where T2D risk variants affect islet regulatory activity^7^, *ZZEF1*, and a recently identified association at *HLA-DRB5^5^.* Heterogeneity in effect direction at specific loci has been observed in other contexts, for example, between T2D and cardiovascular disease and T2D and birth weight^5,26^. We further observed enrichment of nominally associated variants with opposite effects on T1D and T2D in islet regulatory elements, suggesting the potential of a broader mechanistic role for aspects of pancreatic and islet function in opposed risk of T1D and T2D. The specific mechanisms, however, of how *CTRB1, TCF7L2* and other loci encode opposing risk is currently unclear and may involve multiple genes and other cell types.

Another shared mechanism of T1D and T2D pathogenesis is through obesity and insulin resistance. The ‘accelerator’ hypothesis posits that weight gain and insulin resistance exacerbate beta cell stress and T1D progression in a manner similar to T2D pathogenesis^11^. We identified support for this hypothesis through a correlation between increased fasting insulin level and T1D and T2D risk. We also identified enrichment of T1D and T2D variants for adipose and B cell regulatory elements, cell types both involved in insulin resistance. We did not find significant correlation between T1D risk and BMI, or association with large effect obesity loci such as *FTO*. A recent study identified a causal relationship between childhood obesity and T1D risk, supporting a role for adolescent growth in T1D pathogenesis^28^, though we did not observe a genome-wide correlation. There was, however, a positive correlation with male pubertal phenotypes, in line with increased prevalence of T1D in males in early adulthood^41^, and risk variants at the *CENPW* locus were associated with male pubertal growth, height and age of menarche^31,32^. This supports a role for insulin resistance and growth in the shared etiology of T1D and T2D.

We also observed evidence for correlations with other traits, such as between increased T1D and T2D risk and decreased birth weight and increased proinsulin level. Previous studies have reported a correlation between low birth weight and increased T2D risk^26,42^, although the potential link between birth weight and T1D risk is unclear^43^. Furthermore, variants in endoderm regulatory sites were enriched for T1D and T2D association, suggesting potential shared effects on developmental regulatory processes. Proinsulin is an autoantibody in T1D and higher proinsulin level could contribute to increased risk of developing T1D^44^. Conversely, impaired insulin processing is observed in beta cell dysfunction and thus could also represent a consequence of disease progression^45^. Additional studies will be needed to determine causal relationships between proinsulin level or birth weight and diabetes risk and the direction of these relationships.

In total, our findings support shared risk involving variants affecting islet function as well as insulin resistance, growth and development, in the etiology of T1D and T2D. Further studies will help establish the cellular mechanisms of these effects and their role in diabetes pathogenesis.

## Methods

### T1D sample collection

For the type 1 diabetes GWAS, we compiled publicly available genotype-level data for case and control samples from the T1DGC (dbGAP: phs000180.v3.p2), GoKIND/GAIN (dbGAP: phs000018.v2.p1), DCCT-EDIC (dbGAP: phs000086.v3.p1), WTCCC1^46^, and WTCCC2, which were either genotyped on Affymetrix or Illumina platforms (**Table S1**). Because the GoKIND/GAIN dataset contained family trios, we extracted only the proband samples. From the WTCCC1 samples, we used the T1D cohort as cases and the 1958 Birth Cohort (58BC), UK National Blood Service (NBS), and bipolar disorder (BP) cohorts as controls. Unlike a previous study for T1D^47^, we did not include type 2 diabetes or hypertension from WTCCC1 as controls. From the WTCCC2 samples, we used control cohorts from the UK National Blood Service.

### T1D quality control and imputation

We used the recommended individual and variant exclusion lists where available for 58BC, NBS, WTCCC1 T1D and BP. We used phenotype files for GoKIND/GAIN and DCCT-EDIC to exclude samples that were not reported of Caucasian ancestry. We used PLINK^48^ (https://www.cog-genomics.org/plink2) to perform PCA with 1000 Genomes Project (1KGP) samples to identify and remove outliers that did not overlap European 1KGP samples on PC1 and PC2. We used PLINK to calculate identity-by-descent (IBD) values between individuals. Pairs of individuals with at least second-degree relationships (IBD>.2) were pruned in a manner such that only one related individual was retained. For the NBS samples that overlapped between Affymetrix and Illumina platforms, we prioritized the samples genotyped on the Illumina platform. For each cohort, we filtered out variants with less than 95% call rate, less than 1% minor allele frequency (MAF), and extreme Hardy-Weinberg equilibrium values (P<1×10^-5^). We also removed individuals with more than 5% missing genotypes. We then combined cohorts that were genotyped on similar platforms. After filtering steps, the total number of individuals available was 15,043, including 8,967 cases and 6,076 controls (**Table S3**). We imputed 347,083 (Affymetrix) and 500,096 (Illumina) autosomal variants separately into the HRC panel r1.1 using the Michigan Imputation Server^49^, resulting in data for 39,117,105 variants. We excluded variants after imputation that had an imputation quality (r^2^) less than 0.3, leaving 23,385,104 (Affymetrix) and 25,294,976 (Illumina) well-imputed variants.

### T1D genome-wide association and meta-analysis

We used the firth bias-corrected logistic likelihood ratio test as implemented in EPACTS (https://genome.sph.umich.edu/wiki/EPACTS) to test variants for association to T1D separately for Affymetrix and Illumina combined cohorts. We used PLINK to LD prune genotyped variants to create a set of independent variants. We then used PLINK to perform principal component analysis (PCA) and extracted the top 3 principal components (PCs). We used sex and the top 3 PCs as covariates, set a lower MAF threshold of 0.005, and used genotype dosages for association testing. For triallelic SNPs and cases where multiple variants mapped to the same genomic coordinates, we kept the variant with the highest MAF. We then used inverse-variance meta-analysis as implemented in METAL^50^ on association results for 8,720,060 (Affymetrix) and 8,778,018 (Illumina) variants, keeping variants that were tested on both platforms. We further removed genotyped variants that had an empirical R^2^ (ER^2^<.8) for either cohort and all variants in at least moderate LD (r^2^>.5) with these variants. A total of 8,491,085 variants remained for downstream analyses.

To address the potential for misdiagnosed T2D cases in the T1D GWAS, we used phenotype data to remove 278 T1D cases with body-mass index (BMI)>30 from the DCCT and GoKIND/GAIN cohorts. We then re-ran the GWAS meta-analyses using the above methods.

### WTCCC genome-wide association

We collected genotype data for a case cohort of T2D, and control cohorts from NBS and 58BC from the WTCCC1 study^46^. We used sample exclusion lists to remove duplicate, related, or non-Caucasian ancestry samples and SNP exclusion lists to remove poorly genotyped variants. Prior to imputation, we also filtered out variants with less than 95% call rate, less than 1% MAF, and extreme Hardy-Weinberg equilibrium values (P<1e^-5^). We imputed 412,388 genotyped variants from 1,924 T2D case samples and 2,939 control samples together into the HRC panel r1.1 using the Michigan imputation server. After excluding variants with R^2^ < 0.3, we retained 22,520,888 well-imputed variants. We filtered out artifacts by excluding genotyped variants with ER^2^<0.8 and all variants in at least moderate LD (r^2^>.5) with these variants. We used the firth bias-corrected logistic likelihood ratio test as implemented in EPACTS to test variants with MAF > 0.005 for association, using the top 3 PCs as covariates. We finally extracted summary statistics for 1, 173,418 variants in common with the pre-computed European LD score reference panel.

### Genetic enrichment analyses

We tested for enrichment of nominal association and concordance in effects among known T1D and T2D risk loci.

For T2D loci, we collected published credible sets of 49 signals on the Metabochip^19^, 41 additional signals in GoT2D,^8^ and 17 additional signals in DIAGRAM 1000G^4^. We removed all secondary association signals to retain only the primary signal at each locus. For the 93 resulting primary association signals, we then obtained the variant with the highest posterior probability. Where the most likely causal variant was not present in T1D association data, we used the next most likely causal variant. For each variant, we obtained the p-value for T1D association and direction of T1D effect for the T2D risk allele. We tested for enrichment of variants with nominal association (P<.05) by comparing to the expected percentage obtained from sets of matched variants from SNPsnap^51^ using a binomial test.

We then determined concordance in T1D effect direction on T2D variants by calculating the number of variants with same effect direction and applying a binomial test. We further determined the concordance in effect direction in T1D association data in the UK Biobank (ICD10 code E10 from https://sites.google.com/broadinstitute.org/ukbbgwasresults/home) using a binomial test.

For T1D loci, we obtained the variant with the highest posterior probability in fine-mapping of 57 loci described the sections below. Where the top variant was not present in T2D association data we used the next most probable variant. For each variant, we obtained the p-value for T2D association and direction of T2D effect for the T1D risk allele. We tested for enrichment of nominal association (P<.05) by comparing to the expected percentage obtained from sets of matched variants from SNPsnap^51^ using a binomial test.

We then determined concordance in T2D effect direction on T1D variants by calculating the number of variants with same effect direction and applying a binomial test.

### Genetic correlation analyses

We tested for genetic correlation between T1D and T2D, and related glycemic and anthropometric traits, using LD score regression^16,52^.

We collected quantitative trait data for fasting insulin level, fasting glucose level, HOMA-B, HOMA-IR, HbA1C, and proinsulin level from the MAGIC consortium^27,30,53^, body-mass index (BMI) from the GIANT consortium^54^, and pubertal height (12M, 10F), birth weight and childhood obesity from the EGG consortium^26,55^. For the UK Biobank, we obtained summary statistic data of 337k samples using T1D and T2D phenotypes defined from ICD10 codes E10 (T1D) and E11 (T2D) available at sites.google.com/broadinstitute.org/ukbbgwasresults/home. For T2D we obtained data from the GoT2D, HapMap2, and trans-ethnic GWAS studies from the DIAGRAM consortium website.

For each trait, we formatted summary statistics to retain only variants in HapMap3 and correctly orient variant alleles. We then ran LD score regression on the resulting formatted files using default LD scores.

### Genomic enrichment analyses

We considered active enhancer and promoter site annotations for 98 cell types from the Epigenome Roadmap project^21^, along with annotations for coding exons from GENCODE^29^. We used stratified LD-score regression^17^ to identify annotations that were enriched for signal in T1D and T2D association data. Stratified LD-score regression is a multiple regression, where the chi-squared statistics for a trait are regressed on LD-scores computed using variants from each of a set of functional annotations, and the estimated parameters quantify the relative contribution of each annotation to the total heritability.

For the five cell-types with positive enrichment for both T1D and T2D association (pancreatic islets, pancreas, adipose, CD19+ B cells, and CD184+ endoderm), we tested whether these annotations were enriched in variants with shared or opposite effects on T1D and T2D. We identified variants with P<.05 for both T1D and T2D association and in 1000 Genomes phase 3 data. For each of these variants i, we computed z_i,T1D_ = β _i,T1D_ / SE _i,T1D_ and z_i,T2D_ = β_i,T2D_ / SE_i,T2D_. We sorted them by the value of | z_i,T1D_ + z_i,T2D_ | for LD-pruning purposes. After sorting, we pruned these variants using the SNPclip tool of LDlink^54^ using EUR populations, a R^2^>0.1 and MAF>0.01, resulting in 3856 and 2254 independent shared and opposite variants, respectively. We then generated sets of randomized, matched SNPs using SNPsnap^55^. We tested shared and opposite variants for enriched overlap compared to the average overlap across matched variant sets using a one-sided Fisher exact test.

### Fine-mapping of causal variant sets

We used effect and standard error estimates to calculate a Bayes Factor^56^ for each variant. We obtained 58 known loci for T1D from Immunobase and excluded the MHC locus (Table S2). We extracted the previously reported index variants and used PLINK to calculate r^2^ values between 57 index variants and all common variants (MAF>.5) within a 5 MB window as done in a previous study^8^. We defined credible sets of variants for each locus as variants with r^2^>.1 with the index variant. For each locus, we calculated the posterior probability of association (PPA) for each variant by dividing the Bayes Factor for each variant by the sum of Bayes Factors for the entire locus. We then calculated the 99% credible set by taking the set of variants for each locus that added up to 99% PPA. We compared our T1D credible sets to previously published Immunochip credible sets^3^ by extracting 34 common loci between both studies. From the Immunochip study, we extracted only the primary signals. To directly compare p-values, we filtered for variants covered by both studies with non-missing p-values and calculated the Pearson correlation. To identify high probability variants not in Immunochip credible sets, we extracted variants from the 34 loci that were not in the Immunochip primary signal credible set and sorted by PPA.

### Genomic annotations at fine-mapped signals

We considered active regulatory site annotations for cell-types enriched for T1D/T2D association along with annotations for coding exons and UTR regions from GENCODE^29^. For T1D we used fine-mapping data from 57 signals as described above. For T2D we used published fine-mapping data for 93 primary signals from Metabochip, GoT2D and DIAGRAM 1000G studies. At each signal, we calculated a cumulative posterior causal probability (PPA) for each annotation as the sum of PPA values for variants overlapping that annotation. We then assigned T1D/T2D signals to groups based on the highest cumulative PPA value across annotations, considering signals with a cumulative PPA value less than .1 for all annotations as ‘un-annotated’. For each T1D group we then calculated the median association of signals in the group with T2D, and for each T2D group we calculated the median association with T1D.

### Risk signal co-localization

We used a Bayesian co-localization method to determine loci at which T1D and T2D association data showed evidence of a causal variant shared by both traits^57^. At a given locus, the method takes as inputs Bayes Factors of association from two datasets and a specification of the prior probability that each is causal for one or both traits. From these a posterior probability (PP) is computed for each of five hypotheses:

H0: The locus contains no variant causal for either trait

H1: The locus contains a variant causal for trait 1 but none causal for trait 2

H2: The locus contains a variant causal for trait 2 but none for trait 1

H3: The locus contains a variant causal for trait 1 and an independent variant causal for trait 2

H4: The locus contains a variant causal for both H1 and H2.

We used the default prior assumption that all variants at a locus are equally likely to be causal. This model has two important limitations: It assumes each locus has at most one causal variant, and the distinction between H3 and H4 may be confounded by cases of high LD. We considered the prior probability that a variant is associated with T1D or T2D as 1×10^-4^ and the prior probability that a variant is associated with both traits as 1×10^-5^.

We collected 93 T2D loci and 56 T1D loci, of which five have overlapping coordinates (*CENPW, GLIS3, RASGRP1, CTRB1, MTMR3*), for a total of 144 loci (**Supplemental Table 3**). At each locus, we obtained a reported index variant and then extracted all variants in a 500kb window. For each variant, we calculated a Bayes Factor for T1D and T2D separately using the approach of Wakefield^56^. We then applied the co-localization test to compare T1D and T2D Bayes Factors, and considered loci with H4 > .50 as shared. For loci with evidence for a shared risk variant, we then fine-mapped variants causal for the shared signal. For each locus, we multiplied T1D and T2D Bayes Factors at each variant, and then calculated the posterior causal probability (PPA) as the Bayes Factor divided by the sum of all variant Bayes Factors across the locus. We further calculated a cumulative PPA (cPPA) as the sum of PPA values for variants overlapping an annotation at a given locus.

To validate loci with evidence for a shared causal variant we further applied eCAVIAR, a colocalization method capable of modeling multiple causal variants^58^. For each locus, we chose a window of 100 variants on either side of the variant with the strongest combined T1D and T2D evidence. We provided Z-scores of T1D and T2D association together with pairwise LD statistics of European samples in 1000 Genomes Project v3 data for all variants within the window to eCAVIAR using default settings. For each variant in the window, eCAVIAR computed a colocalization posterior probability (CLPP), the probability that the variant is causal for the local signal in both traits. We considered loci to be co-localized using this approach with at least one variant with CLPP > 0.01 as recommended in the original study.

For quantitative trait association at shared risk variants, we obtained the most likely causal variant from combined T1D and T2D evidence. We extracted summary statistics for each trait and calculated a signed Z-score for the risk allele using effect size and standard error estimates.

### Islet ATAC-seq and chromatin states

We utilized ATAC-seq data generated from four primary pancreatic islet samples as described in a separate study^59^. For each sample, we trimmed adaptor sequences from the reads with trim_galore (https://github.com/FelixKrueger/TrimGalore). The resulting sequences were aligned to sex-specific hg19 reference genomes using bwa mem^60^. We filtered reads to retain those in proper pairs and with mapping quality score greater than 30. We then removed duplicate and non-autosomal reads. We called sites individually for each sample with MACS2^61^ at a q-value threshold of .05 with the following options “—no-model”, “—shift −100”, “—extsize 200”. We removed sites that overlapped genomic regions blacklisted by the ENCODE consortium^20^. We merged sites from these 4 samples and two previously generated in islets^33^ with bedtools^62^ to obtain a comprehensive set of ATAC-seq peaks in human islets.

We used islet chromatin states described separately^34^. In brief, we used previously published data^6,35^ from ChIP-seq assays generated in islets and for which there was matching input sequence from the same sample. For each assay and input, we aligned reads to the human genome hg19 using bwa samse and bwa aln^60^ with a flag to trim reads at a quality threshold of less than 15. We converted the alignments to bam format and sorted the bam files. We then removed duplicate reads, and further filtered reads that had a mapping quality score below 30. Sequence data from the same assay in the same sample were then pooled. We defined chromatin states from ChIP-seq data using ChromHMM^63^ with a 9 state model. We assigned the resulting states names based on the resulting patterns.

### ATAC-seq footprint analysis

To identify haplotype-aware motifs within ATAC-seq footprints overlapping accessible chromatin sites, we searched accessible chromatin sites from four ATAC-seq samples for instances of motifs from JASPAR, SELEX, ENCODE and *de novo* motifs identified in our data^64^. We used vcf2diploid^65^ (https://github.com/abyzovlab/vcf2diploid) to create individual-specific diploid genomes by mapping our phased, imputed genotypes onto hg19 using only SNPs and ignoring indels. Then, we used fimo^66^ to scan the personalized genomes for our compiled database of motifs, limiting the sequences scanned to those derived from islet accessible chromatin. For fimo, we used the default parameters for p-value threshold (1×10^-4^) and a background GC content of 40.9% based on hg19.

CENTIPEDE^67^ was used to discover footprint sites for each motif, using the discovered motif instances within ATAC-seq peaks. For each motif, we used the make_cut_matrix utility from atactk (https://github.com/ParkerLab/atactk) to calculate a cut-site matrix that contained counts of the number of Tn5 integrations within a window defined by ±100 bp from each motif occurrence for both forward and reverse strands. This cut-site matrix was provided as input to CENTIPEDE along with regions for each motif occurrence to model the posterior probability that a given motif occurrence was bound by a TF. We defined footprints for a given motif as regions that had a posterior probability ≥ 0.99. We combined footprints from our samples with a previously published set of footprints in pancreatic islets^33^.

We further identified variants predicted to disrupt each footprint^4^. We calculated the entropy score for a variant position in a footprint using the position frequency matrix for each motif. For each base at a given position bp and the frequency of the base at that position f, we calculated the entropy as:

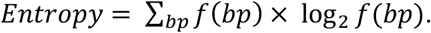

A footprint was considered disrupted if a variant fell in a conserved position in the motif (Entropy<1.0).

### Luciferase reporter assays

To test for allelic differences in enhancer activity at rs4237150, rs10116772 and rs8056814, we cloned sequences containing the alt or ref allele in forward and reverse orientation upstream of the minimal promoter of firefly luciferase vector pGL4.23 (Promega) using KpnI and SacI restriction sites.

Primer sequences were:

rs4237150

Fwd: TTACGCGGTACCACACACTTCTGTAAATCAGGTCAG, TCATAGGAGCTCGAAGCAGTTTGTTTGCTGGC

Rev: TTACGCGAGCTCACACACTTCTGTAAATCAGGTCAG, TCATAGGGTACCGAAGCAGTTTGTTTGCTGGC

rs6476839

Fwd: GTCGGTACCTCGCAATTCAATCAAGGACA, GCTGAGCTCCAGGCACATGTTTGCACTTT

Rev: GTCGAGTCGTCGCAATTCAATCAAGGACA, GCTGGTACCCAGGCACATGTTTGCACTTT

rs10116772+rs10814915

Fwd: GTCGGTACCTTCATTAATGCCGCCTTTTC, GCTGAGCTCTGAATTGCGAAATGTGCTTC

Rev: GTCGAGTCGTTCATTAATGCCGCCTTTTC, GCTGGTACCTGAATTGCGAAATGTGCTTC

rs8056814

Fwd: TAAGCAGGTACCTGGGTGACAGAGTGAGACTCC, TGCTTAGAGCTCGGTGTTTCCGCCTAACACTG

Rev:TAAGCAGAGCTCTGGGTGACAGAGTGAGACTCC,TGCTTAGGTACCGGTGTTTCCGCCTAACACTG

MIN6 beta cells were seeded into 6 (or 12)-well trays at 1 million cells per well. At 80% confluency, cells were co-transfected with 400ng of the experimental firefly luciferase vector pGL4.23 containing the alt or ref allele in either orientation or an empty vector and 50ng of the vector pRL-SV40 (Promega) using the Lipofectamine 3000 reagent. All transfections were done in triplicate. Cells were lysed 48 hours after transfection and assayed for Firefly and Renilla luciferase activities using the Dual-Luciferase Reporter system (Promega). Firefly activity was normalized to Renilla activity and compared to the empty vector and normalized results were expressed as fold change compared to empty vector control per allele. A two-sided t-test was used to compare the luciferase activity between the two alleles in each orientation.

### Allelic imbalance analysis

We collected ChIP-seq data from assays in primary islet cells from multiple sources^6,35,68–71^. We aligned sequence data using bwa samse^60^, filtered out mitochondrial reads, and removed duplicates using Picard software. For each sample we applied QuASAR^72^ to obtain estimated genotypes. A total of 6 samples were determined to be heterozygous at rs4237150 with probability of being homozygous < 10^-4^. For these samples we also inferred heterozygosity at rs10116772, due to high linkage and by imputation into 1000 Genomes v3 variants via the Michigan Imputation Server^49^. Across these 6 samples, a total of 8 datasets had more than 5 reads overlapping rs4237150 – FOXA2 (1), H3K27ac (3), PDX1 (2), NKX6-1 (2). We applied WASP^73^ to each dataset to correct for reference mapping bias. We then pooled read counts for risk and protective alleles at rs4237150 and rs10116772 and applied a two-sided binomial test for allelic imbalance.

### Author Contributions

K.J.G. designed the study; K.J.G, A.J.A. and J.C. wrote the manuscript and performed genetic and genomic analyses; M.O. and N.K. performed experiments and contributed to analyses.

## Acknowledgements

This work in this manuscript supported in part by NIH/NIDDK award DK112155 and ADA award 1-17-JDF-027 to KJG. <u>GoKinD</u>: The Genetics of Kidneys in Diabetes (GoKinD) Study was conducted by the GoKinD Investigators and supported by the Juvenile Diabetes Research Foundation, the CDC, and the Special Statutory Funding Program for Type 1 Diabetes Research administered by the National Institute of Diabetes and Digestive and Kidney Diseases (NIDDK). The data from the GoKinD study were supplied by the NIDDK Central Repositories. <u>DCCT/EDIC</u>: The Diabetes Control and Complications Trial (DCCT) and its follow-up the Epidemiology of Diabetes Interventions and Complications (EDIC) study were conducted by the DCCT/EDIC Research Group and supported by National Institute of Health grants and contracts and by the General Clinical Research Center Program, NCRR. The data from the DCCT/EDIC study were supplied by the NIDDK Central Repositories. <u>T1DGC</u>: This research utilizes resources provided by the Type 1 Diabetes Genetics Consortium (T1DGC), a collaborative clinical study sponsored by the National Institute of Diabetes and Digestive and Kidney Diseases (NIDDK), National Institute of Allergy and Infectious Diseases (NIAID), National Human Genome Research Institute (NHGRI), National Institute of Child Health and Human Development (NICHD), and the Juvenile Diabetes Research Foundation International (JDRF) and supported by U01 DK062418. The UK case series collection was additionally funded by the JDRF and Wellcome Trust and the National Institute for Health Research Cambridge Biomedical Centre, at the Cambridge Institute for Medical Research, UK (CIMR), which is in receipt of a Wellcome Trust Strategic Award (079895). The data from the T1DGC study were supplied by the NIDDK Central Repositories. <u>WTCCC</u>: This study makes use of data generated by the Wellcome Trust Case Control Consortium. A full list of the investigators who contributed to the generation of the data is available from www.wtccc.org.uk. Funding for the project was provided by the Wellcome Trust under award 076113. This manuscript was not prepared in collaboration with investigators of these studies and does not necessarily reflect the opinions or views of the WTCCC, GoKinD, DCCT/EDIC or T1 DGC studies or study groups, the NIDDK Central Repositories, the NIH, or the study sponsors.

## Data availability

Summary data will be available at http://www.gaultonlab.org/pages/Aylward_T1D_T2D

## Supplemental Figures

**Supplemental Figure 1.**
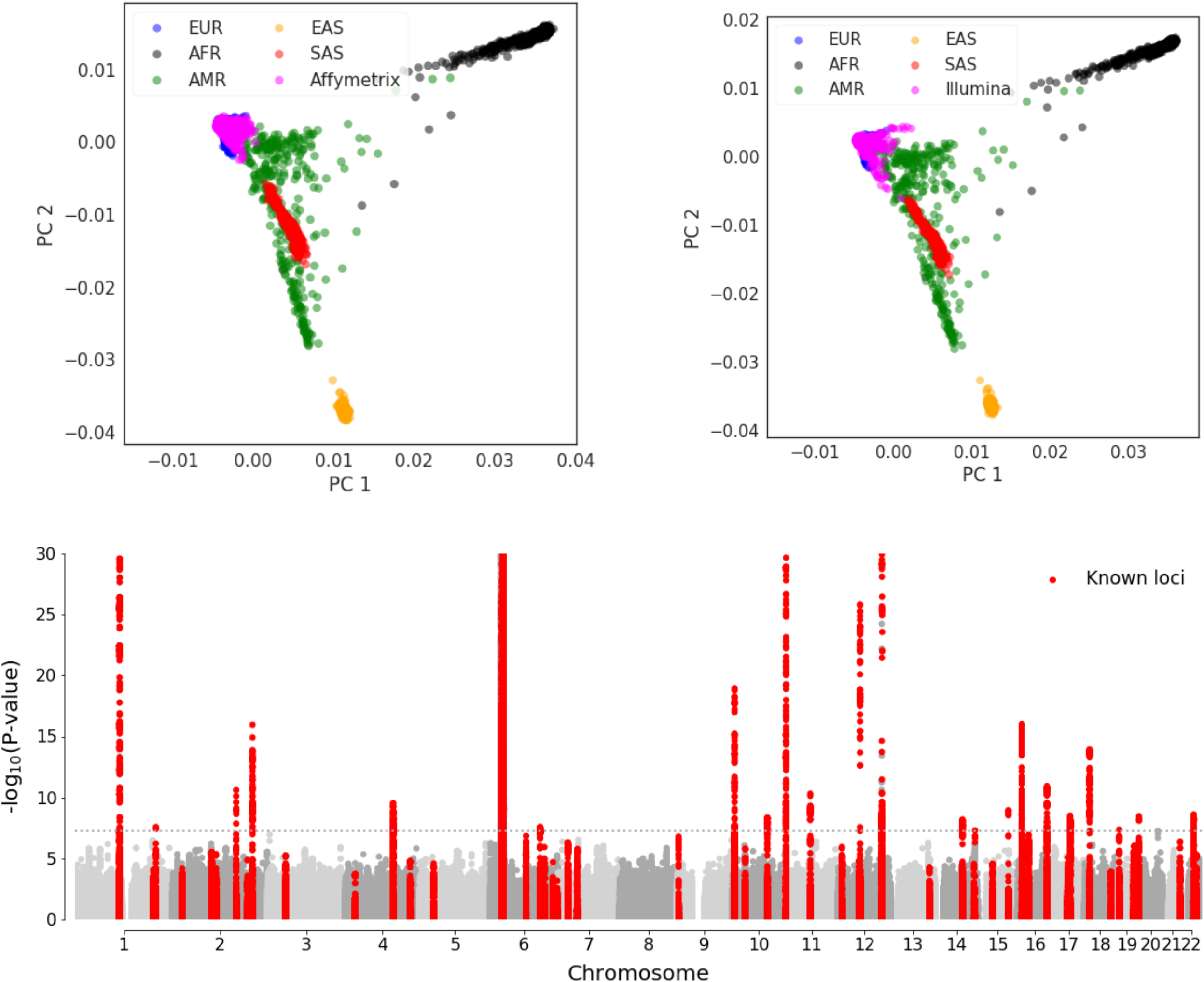
Genome-wide association study of T1D case and control samples. (A) Principal component plots showing the ancestry of samples genotyped on Affymetrix and Illumina arrays as compared to the super populations of the 1000 Genomes Project after QC measures. EUR = European, AFR = African, AMR = Americas, EAS = East Asian, and SAS = South Asian. (B) Manhattan plot plotting chromosomal positions (hg19) and the negative log10-P-values, with known T1D loci highlighted in red.

**Supplemental Figure 2.**
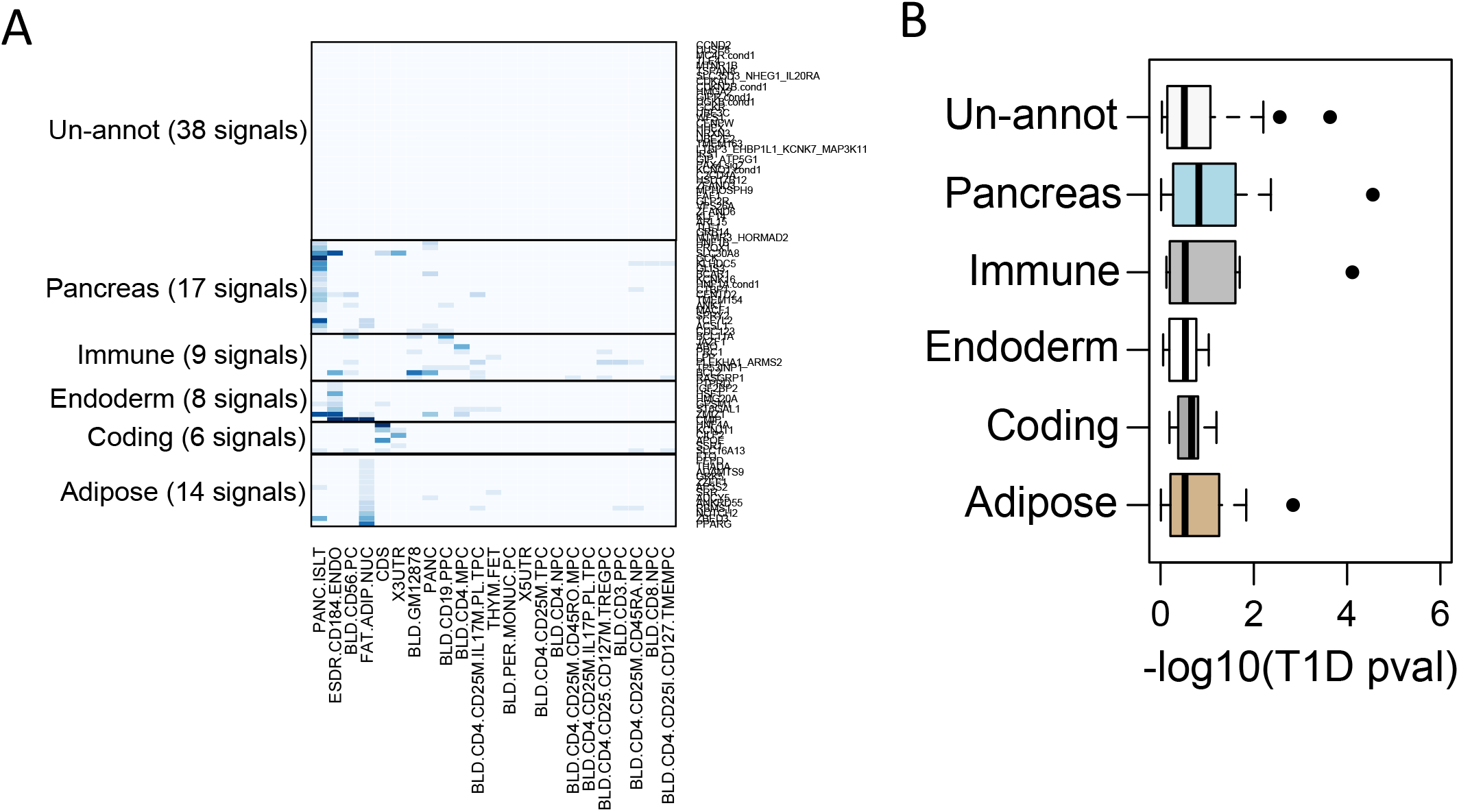
Genomic annotations and T1D association and fine-mapped T2D loci. (A) Cumulative PPA values of 93 primary T2D signals in cell-type regulatory site and coding annotations. T2D signals mapped into six groups including pancreatic regulatory sites (21 signals), adipose regulatory sites (15 signals), and coding exons (4 signals) in addition to unannotated signals. (B) T2D signals within different groups had distinct patterns of association with T1D, where T2D pancreas signals had the strongest T1D association.

**Supplemental Figure 3.**
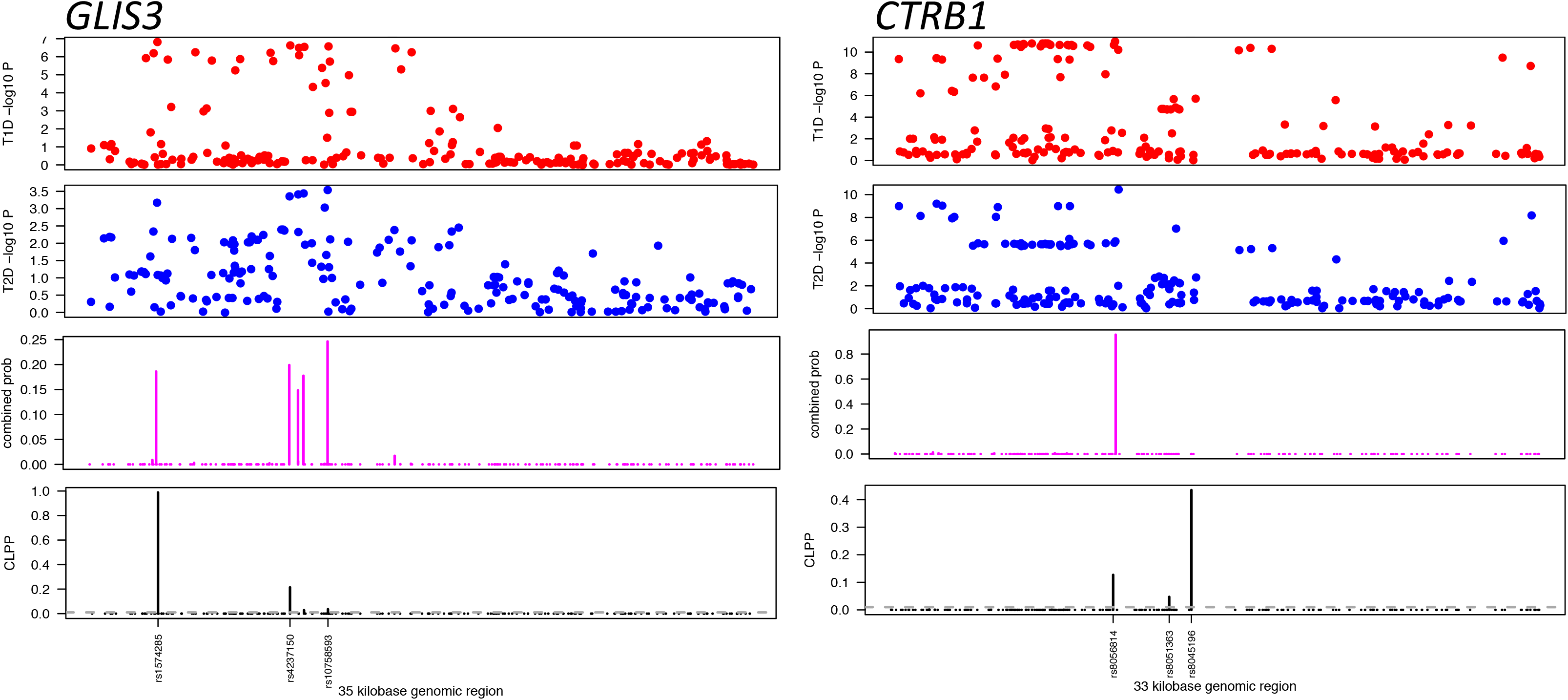
Shared T1D and T2D signals at the *GLIS3* and *CTRB1* loci. (top) P-values of variant associations with T1D (red) and T2D (blue) at the *GLIS3* and *CTRB1* loci. Causal probability of variants at the shared *GLIS3* and *CTRB1* signals by (middle) combining T1D and T2D evidence in Bayesian fine-mapping, and (bottom) modeling shared causal variants using eCAVIAR. Variants at each signal have high causal probabilities in both analyses.

**Supplemental Figure 4.**
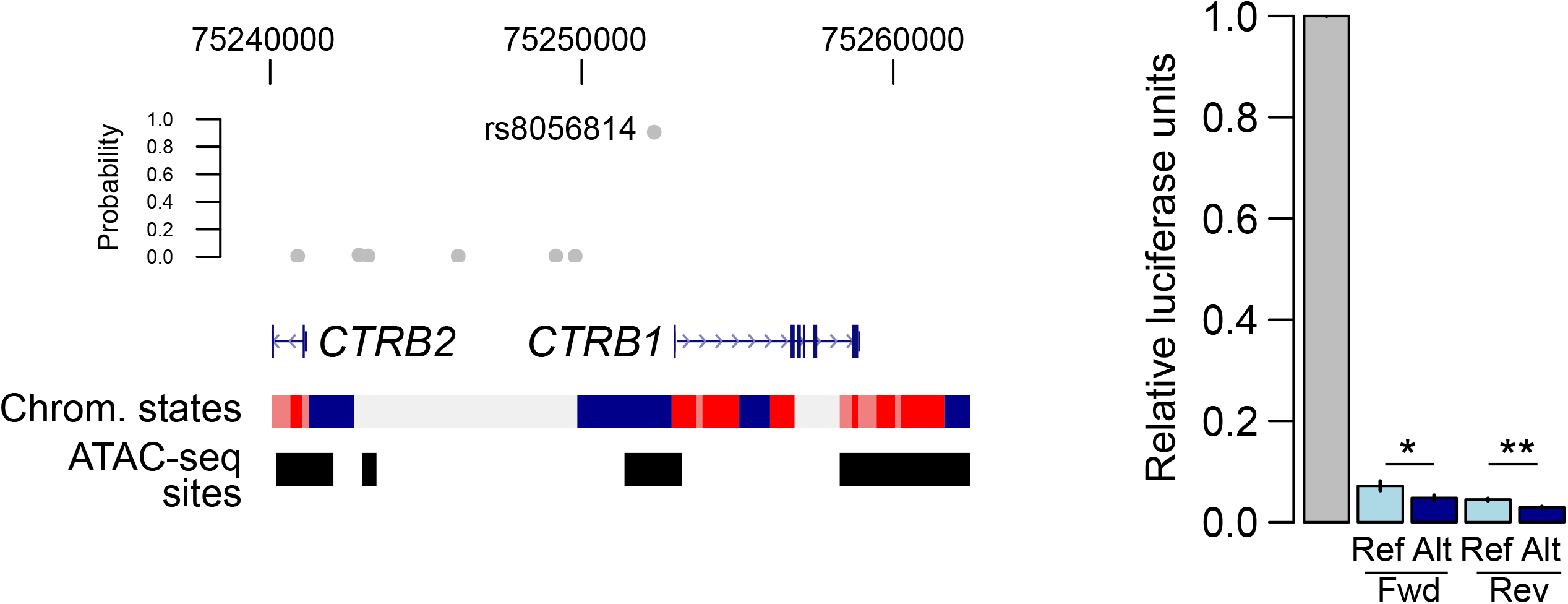
Allelic imbalance in islet regulatory activity at *GLIS3.* Read counts in samples heterozygote for rs4237150 and rs10116772 in pancreatic islet FOXA1, NKX6.1, PDX1 and H3K27ac ChIP-seq assays (risk allele counts = light grey, protective allele = dark grey). The risk allele had increased read counts in all assays. P-values for binomial tests are listed below each assay.

**Supplemental Figure 5.**
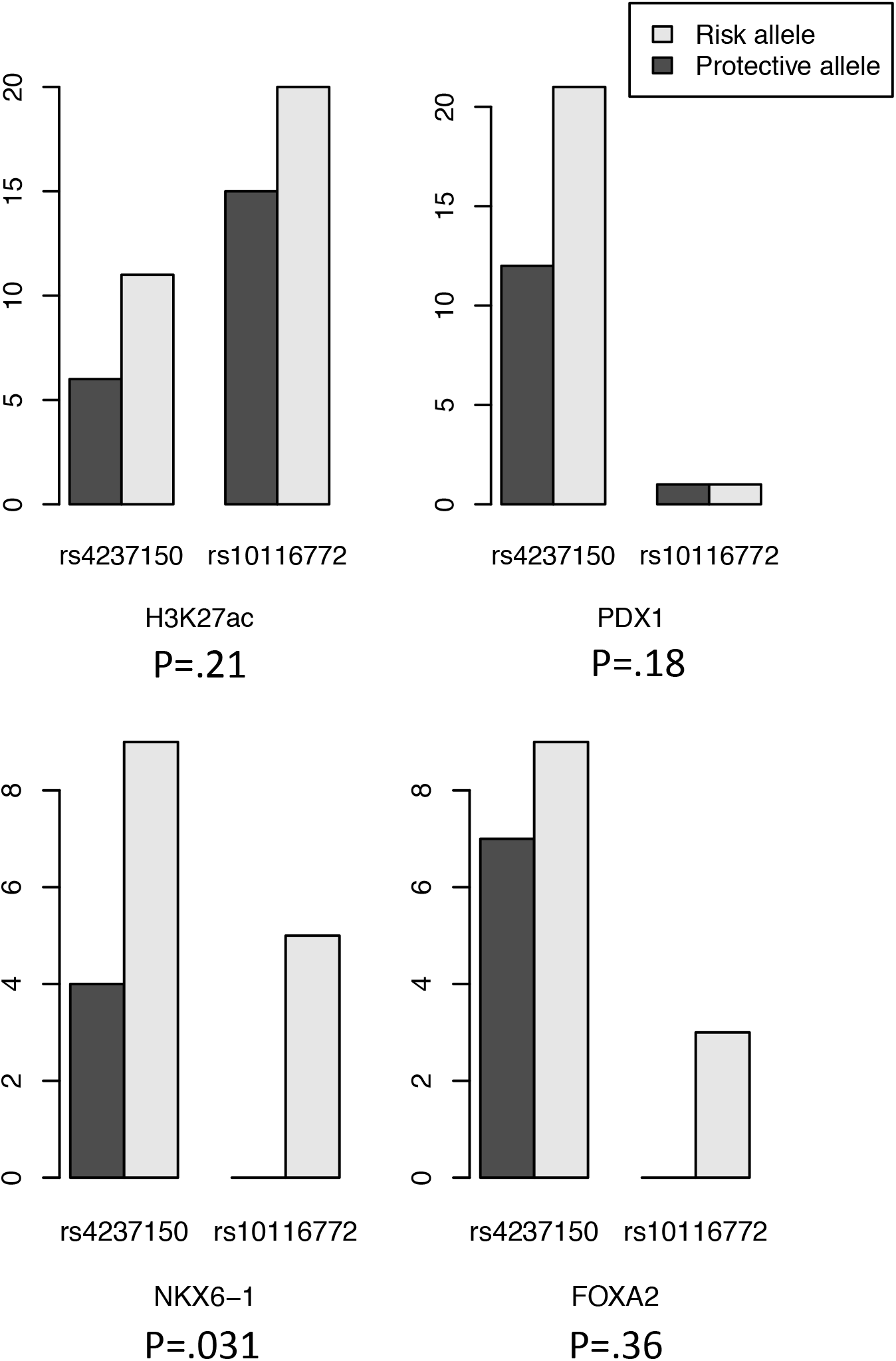
Shared variant at *CTRB1* affects islet regulatory activity. (A) Plot of candidate causal variants at the shared *CTRB1* signal. Variant rs8056814 has a high probability (PPA=.90) of being causal for T1D and T2D risk, and maps in an islet accessible chromatin site and an islet active enhancer upstream of *CTRB1.* (B) Luciferase reporter assay of sequence surrounding rs8056814 alleles in the islet cell line MIN6. All constructs had reduced activity compared to the empty vector. The T2D risk allele of rs8056814 has increased activity compared to the T2D protective allele. Values are fold change and SD. (N=3; *P<.05, **P<.001).

